# Hearing Lips and Seeing Voices After Fifty Years: A Large-Scale McGurk Illusion Dataset for Audiovisual Speech Research

**DOI:** 10.64898/2026.06.09.731046

**Authors:** Zhengye Wang, Gantang Li, Yating Yu, Junyi Wu, Zenghui Yu, Yang Meng, Suiping Wang, Chenjie Dong

## Abstract

Efficient face-to-face communication relies on the integration of auditory speech and visual articulatory signals. Over the past five decades, the McGurk illusion has been widely used as an index of audiovisual speech integration. However, substantial variabilities in susceptibility to the illusion across participants and speakers limit its reliability as a stable measure of audiovisual integration ability. Here, we introduce the McGurk illusion dataset (MID), which, to our knowledge, is the largest publicly available McGurk stimulus dataset to date. The MID comprises auditory (N = 400), visual (N = 400), and audiovisual (N = 640) speech stimuli generated from 80 Mandarin speakers and validated through behavioral judgments across 360,900 trials. Using this dataset, we characterized the acoustic and facial articulatory properties of McGurk stimuli, replicated substantial inter-participant and inter-speaker variabilities in illusion susceptibility, and revealed the associations between variations in McGurk illusion rate and the variations in unisensory perception, audiovisual correspondences, and speakers’ characteristics. Furthermore, the stimulus set enabled systematic comparisons of the reliability of different McGurk illusion-based indices of audiovisual speech integration. Overall, the MID not only provides a standardized resource for investigating audiovisual speech integration and its alterations across populations, but also supports research on speaker normalization, lip-reading, and speech perception.

## 1. Introduction

Efficient face-to-face communication relies on the integration of auditory speech and visual articulatory signals (Young et al., 2020). Audiovisual speech integration substantially improves communication in challenging listening environments (Bernstein et al., 2004; Oh et al., 2022), but it can also give rise to perceptual illusions under conditions of audiovisual conflict, as exemplified by the McGurk illusion (Alsius et al., 2018; McGurk & MacDonald, 1976). In a typical McGurk stimulus, an auditory syllable (e.g., /ba/) is paired with an incongruent visual articulation (e.g., /ga/), often leading listeners to perceive an illusory syllable such as /da/ (McGurk & MacDonald, 1976).

Over the past five decades, the McGurk illusion has been widely used as an index of audiovisual speech integration because of its intuitiveness and ease of implementation (Alsius et al., 2018; Wallace et al., 2020). In neurotypical populations, susceptibility to the illusion has been associated with genetic factors (Feng et al., 2019), gender differences (Irwin et al., 2006), and cultural backgrounds (Magnotti et al., 2025; Meronen et al., 2013; Mohamed et al., 2024) in audiovisual integration. Altered susceptibility has also been reported in several clinical populations, including individuals with autism (Stevenson et al., 2014), schizophrenia (Roa Romero et al., 2016), dyslexia (Bastien-Toniazzo et al., 2010), and hearing deficits (Stropahl et al., 2017). More recently, the McGurk illusion has also been used to investigate the speech processing of multimodal artificial intelligence (Ma et al., 2026).

Despite its widespread use, studies have reported substantial variability in susceptibility to the McGurk illusion among neurotypical participants, with illusion rates ranging from 0% to 100% even when participants were tested with the same stimulus (Basu Mallick et al., 2015; Dong et al., 2024). This inter-participant variability was partly predicted by perceptual precision on the corresponding unisensory syllables (Brown et al., 2018; Dong et al., 2025; Strand et al., 2014). Moreover, susceptibility to the McGurk illusion shows relatively weak associations with other measures of audiovisual speech perception, such as speech-in-noise recognition (Magnotti et al., 2020; Van Engen et al., 2017). These findings raise concerns about the reliability and generalizability of the McGurk illusion as a measure of audiovisual integration ability (Getz & Toscano, 2021; Van Engen et al., 2022).

In addition to variability across participants, studies have revealed substantial speaker-dependent variability (i.e., inter-speaker variability) in susceptibility to the McGurk illusion (Basu Mallick et al., 2015; Magnotti et al., 2024). For example, Dong and colleagues reported that when participants were tested with McGurk stimuli from 20 speakers, illusion rates within a single participant ranged from 0% to 100% depending on the speaker (Dong et al., 2025). Furthermore, susceptibility to a given McGurk stimulus was partly predicted by perceptual performance for the corresponding unisensory syllables produced by that speaker. These findings suggest that susceptibility to the McGurk illusion is strongly influenced by speaker-specific acoustic and visual articulatory properties.

Intriguingly, despite substantial variability across participants and speakers, susceptibility to a given McGurk stimulus appears to be relatively stable within individuals (Dong et al., 2025) and remains consistent over time (Basu Mallick et al., 2015). Group differences in illusion susceptibility also persist after accounting for unisensory perceptual performance (Bebko et al., 2014). In addition, McGurk illusion rates, indexed by the proportion of illusory “da” responses, were associated with accuracy on audiovisual congruent /da/ syllables at both behavioral and neural levels (Dong et al., 2024). These findings suggest that susceptibility to the McGurk illusion reflects stable but speaker-dependent mechanisms of audiovisual speech perception, highlighting the importance of systematically characterizing McGurk stimuli across speakers.

However, only a limited number of studies have used McGurk stimuli from more than ten speakers (Table 1). Given the substantial variability in McGurk illusion susceptibility across speakers, the heterogeneity of stimuli across laboratories and the limited availability of publicly accessible stimulus sets have hindered comparability, reproducibility, and cumulative progress in audiovisual speech research (Alsius et al., 2018).

**Table 1.** Previous studies of the McGurk illusion incorporating more than five speakers.

| Author (year) | Number of speakers | Syllables |
| --- | --- | --- |
| Dong et al. (2025) | Exp 1: 4 (2 females, 2 males);<br>Exp 2: 3 (1 females, 2 males);<br>Exp 3: 20 (10 females, 10 males) | /ba/, /da/, and /ga/ |
| Magnotti et al. (2025) | 8 (3 females, 5 males) | /ba/, /ga/, /pa/, and /ka/ |
| Magnotti et al. (2024) | 15 (McGurk stimuli/talkers) | /ba/, /ga/, /pa/, and /ka/ |
| Tiippana et al. (2023) | 8 (4 females, 4 males) | /ka/, /pa/, and /ta/ |
| Van Engen et al. (2017) | 8 (4 females, 4 males) | /ba/ and /ga/ |
| Stropahl et al. (2017) | 8 (4 females, 4 males) | /ba/, /da/, /ga/, /ka/, /ma/, /na/, /pa/, and /ta/ |
| Basu Mallick et al. (2015) | 4 females, 4 males | /ba/, /da/, and /ga/ |
| Munhall et al. (1996) | Exp 1: 1 female;<br>Exp 2: 3 females;<br>Exp 3: 2 females | /ba/, /ga/, and /igi/ |

Exp = experiment. Gender distributions of speakers in Magnotti et al. (2024) are omitted, as this demographic information was not specified in the original reports.

To address this gap, we created the largest publicly available McGurk stimulus dataset to date, comprising stimuli from eighty Mandarin speakers. The dataset was acquired under standardized recording conditions and validated through extensive behavioral testing, yielding 360,900 trials in total. Using this dataset, we characterized acoustic and visual articulatory signal of different McGurk syllables (i.e., /ba/, /da/, and /ga/), replicated and extended key findings from previous studies, and systematically compared the reliability of different McGurk-based indices of audiovisual speech integration.

**Fig. 1.**
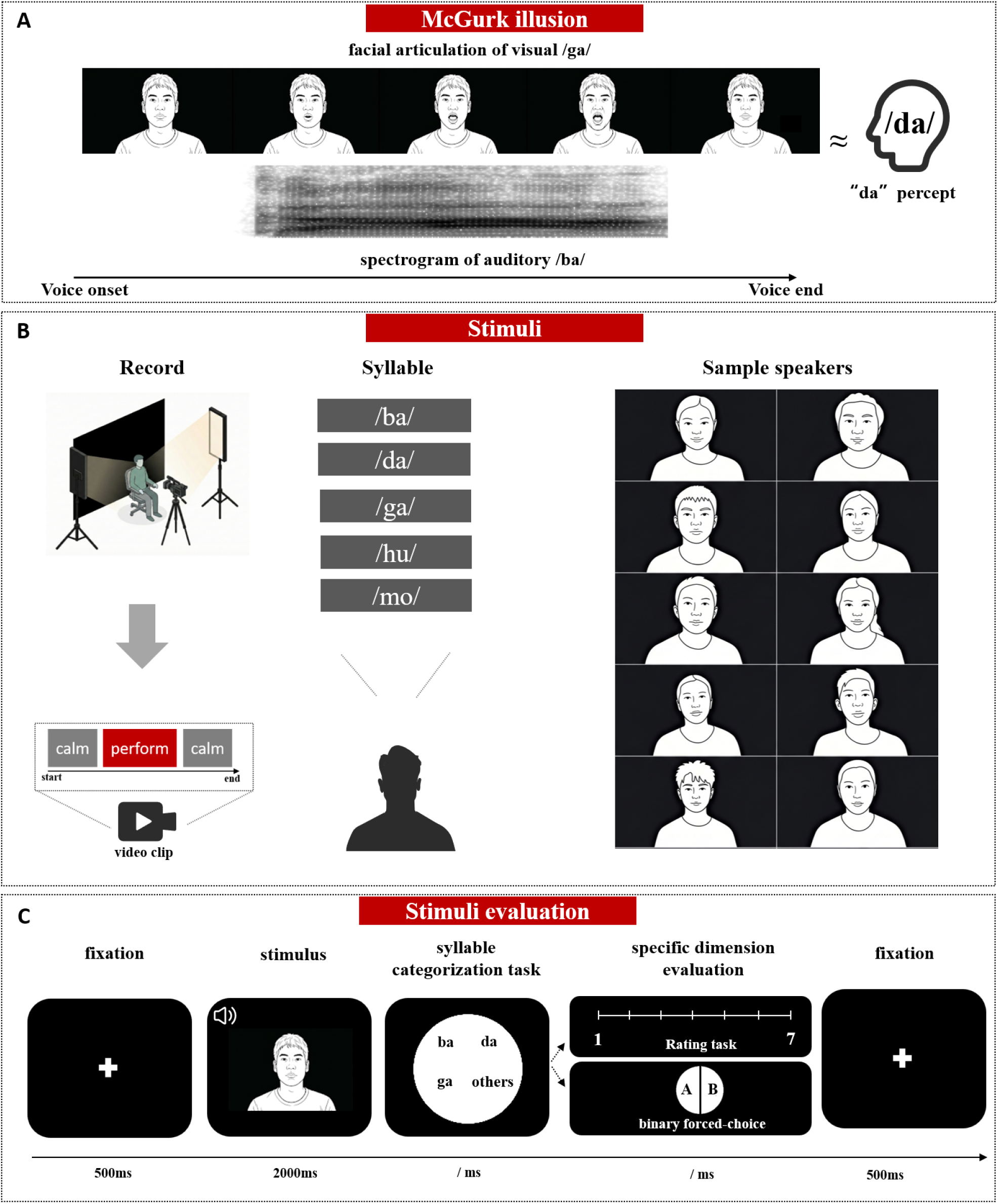
Stimuli and procedures. (A) Demonstration of the McGurk illusion. (B) Stimulus generation and dataset composition; the recording procedure (left), recorded syllables (middle), and exemplar speakers (right). (C) Experimental procedure of the audiovisual judgement task. The photos used in this figure were recordings of lab members who consented to the use of their images in the manuscript.

## 2. Methods

### 2.1 Participants

A total of one hundred and thirty-seven Chinese adults participated in the study. One hundred and seven participants served as speakers of the stimuli (47 women; mean age = 20.88, SD = 2.53), and thirty participants served as perceivers in the behavior judgement (17 women; mean age = 20.43, SD = 2.72). All participants reported normal or corrected-to-normal vision and no history of auditory or neurological disorders.

### 2.2 Materials

#### 2.2.1 Types of stimuli

The dataset comprised five stimulus types: auditory-only (A), visual-only (V), audiovisual congruent (AVc), audiovisual incongruent McGurk (AVm), and audiovisual incongruent non-McGurk stimuli (AVi). Auditory-only stimuli were voice of three targeted McGurk related syllables /ba/ (A_ba_), /da/ (A_da_), /ga/ (A_ga_), and two filler syllables /mo/ (A_mo_)/ and /hu/ (Ahu). Filler syllables were used to reduce the effect of sensory adaptation during behavior judgement. Visual-only stimuli were facial articulation of syllable /ba/ (V_ba_), /da/ (V_da_), /ga/ (V_ga_), /mo/ (V_mo_), and /hu/ (V_hu_).

Audiovisual congruent stimuli were the combination of auditory and visual syllables, including /ba/ (AVc_ba_), /da/ (AVc_da_), /ga/ (AVc_ga_). Audiovisual incongruent McGurk stimuli consisted of the voice of /ba/ pared with the facial articulation of /ga/ (A_ba_+V_ga_ or AV_m_). Audiovisual incongruent nan-McGurk stimuli (AVi) consisted of the voice of /ga/ pared with the facial articulation of /ba/ (A_ga_V_ba_), the voice of /da/ pared with the facial articulation of /ba/ (A_da_V_ba_), the voice of /mo/ pared with the facial articulation of /hu/ (A_mo_V_hu_), and the voice of /hu/ pared with the facial articulation of /mo/(AhuV_mo_).

#### 2.2.2 Stimuli recording, screening, and editing

##### Recording

Stimuli recordings were conducted in a sound-attenuated studio (< 25 dB). Speakers were seated against a black backdrop and illuminated by two 19-inch fill lights positioned at 45° angles. Each syllable was articulated at least three times to ensure the quality of speech. Video and audio were captured with a Sony PXW-Z90V camcorder positioned frontally (0° azimuth, ≤ 5° vertical deviation) to approximate a face-to-face viewing perspective.

##### Screening

One thousand six hundred five raw video clips were recorded from 107 speakers, each pronouncing 5 syllables (i.e., /ba/, /da/, /ga/, /mo/ and /hu/), repeated 3 times. Among these, recordings from twenty-seven speakers were excluded according to the following exclusion criteria: (i) low articulatory clarity, such as incomplete syllable production; (ii) unstable head position or excessive movement during articulation; (iii) visible eye blinking or other distracting facial behaviors; or (iv) poor audio quality, including clipping, background noise, or inconsistent loudness. Raw video clips of the remaining eighty speakers were imported into Adobe Premiere Pro 2023, and the optimal clip with high articulatory clarity, head stability, and no eye blinking was selected from three repetitions for further editing, yielding four hundred valid clips.

##### Editing

Selected clips were trimmed to preserve the full visible articulatory cycle, from initial lip closure through maximal oral opening and back to closure. To standardize facial articulatory signals, each clip underwent a two-step adjustment: (i) the footage was digitally magnified by 200%, and (ii) the frame was spatially aligned so that the philtrum was centered at the origin (x = 0, y = 0). To ensure consistent loudness across auditory stimuli, auditory tracks underwent root-mean-square (RMS) amplitude normalization. The RMS intensity was calculated over the core vocalic segment (the steady vowel portion), and each track was linearly scaled to a standardized target level of -26 dBFS (±0.5 dB). The edited video and audio clips were combined to generate the audiovisual incongruent stimuli. Audio onsets in both audiovisual McGurk and audiovisual incongruent non-McGurk stimuli were aligned to the original congruent audiovisual recordings to ensure temporal congruency. All stimuli were approximately 2 seconds, with 1920 × 1080 resolution and 25 frames per second. Audio tracks were sampled at 48 kHz and presented at 70 dB SPL through noise-attenuating headphones.

### 2.3 Stimuli evaluation

The stimuli evaluation comprised three sessions using a four-alternative forced-choice (4AFC) syllable categorization task: an auditory-only session, a visual-only session, and an audiovisual session. While the trial structure remained consistent across sessions, the stimulus modality and the number of required judgements varied. In the unisensory sessions, participants completed a single judgement (syllable categorization) per trial; in the audiovisual session, they completed two sequential judgements (syllable categorization and characteristic evaluation). Each trial began with a 500 ms fixation, followed by a 2,000 ms stimulus, and a syllable categorization with a 4AFC response screen (unlimited duration). Choice options were presented on a circular interface divided into four equal sectors, with labels “ba,” “da,” “ga,” and “other” presented in randomized sectors on each trial. In the auditory and audiovisual sessions, participants were required to report the syllable they heard by clicking mouse, and in the visual session, they were required to report the syllable they saw.

In the audiovisual session, the 4AFC response screen was followed by a characteristic evaluation screen. On this screen, participants were required to evaluate one of five characteristics of the stimulus randomly presented in each trial: (i) the audiovisual congruency and (ii) the audiovisual synchrony, tested with two-alternative choice options (congruent vs. incongruent, synchrony vs. asynchrony) presented randomly on the left or right of fixation, (iii) the attractiveness of the speaker, (iv) articulatory clarity of the speaker, and (v) similarity between the speaker’s and their own articulation (from now on self-speaker similarity), each assessed on a 7-point Likert scale. Attractiveness was rated from 1 (very unattractive) to 7 (very attractive), articulatory clarity from 1 (very unclear articulation) to 7 (very clear articulation), and self-speaker similarity from 1 (completely unlike one’s own articulatory habits) to 7 (very similar to one’s own articulatory habits). Each characteristic of each syllable produced by each speaker will be assessed twice across the whole task.

The auditory and visual sessions each included 3,200 trials. In each session, /ba/, /da/, and /ga/ stimuli were each repeated 10 times per speaker, whereas /mo/ and /hu/ stimuli were each repeated 5 times per speaker, yielding equal opportunity for the four response categories (“ba,” “da,” “ga,” and “other”). The audiovisual session included 5,630 trials: 2,400 audiovisual congruent trials (10 repetitions per syllable per speaker), 800 audiovisual incongruent McGurk trials (A_ba_V_ga:_ 10 repetitions per speaker), 2,400 audiovisual incongruent non-McGurk trials (A_ga_V_ba_ and A_da_V_ba_: 10 repetitions per speaker; A_mo_V_hu_ and A_hu_V_mo_: 5 repetitions per speaker), and 30 audiovisual asynchronous and incongruent filler trials. These filler trials were generated from AVm and AVi stimuli produced by six speakers (S001–S006), with visual onset preceding auditory onset by 400 ms, and were included to ensure that participants encountered true asynchronous and incongruent audiovisual stimuli during the task.

Each participant completed all the sessions over 10 days, yielding 12,030 trials in total (approximately 25 hours). Session order was counterbalanced by day: auditory-visual-audiovisual on odd days and visual–auditory–audiovisual on even days. To reduce fatigue and practice effects, participants received a unique pseudo-randomized schedule that balanced daily workload. Testing was programmed in PsychoPy (2021.1.2.3), and conducted in a quiet laboratory, with participants seated 60 cm from a 1,920 × 1,080 monitor.

### 2.4 Data analysis

Before conducting formal analyses, we applied a response-time exclusion procedure following Dong et al. (2025). Specifically, trials with response times exceeding three standard deviations from the relevant mean were removed. The total exclusion rates were 1.5% for syllable categorization, 2.2% for audiovisual congruency, 2.1% for audiovisual synchrony, 2.0% for attractiveness, 2.1% for articulatory clarity, and 2.1% for self-speaker similarity judgements. All analyses were implemented using Python (Version 3.12) and R (Version 4.3.1).

#### 2.4.1 Physical properties of the McGurk stimuli

To characterize the physical properties of the McGurk stimuli (auditory /ba/, visual /ga/, and illusory /da/), we extracted acoustic and facial articulatory signal from the audio and video recordings of each speaker. For acoustic signal, the first (F1), second (F2), and third (F3) formant frequencies of the syllables A_ba_, A_da_, A_ga_ were calculated using the Parselmouth interface to Praat. For facial articulatory signal, the normalized mean mouth area and mouth-movement velocity for syllables V_ba_, V_da_, V_ga_ were calculated using MediaPipe Face Mesh.

To examine differences in these physical properties across syllables, we conducted non-parametric Friedman tests across syllables. Post-hoc pairwise comparisons were performed using Wilcoxon signed-rank tests with Bonferroni corrections. Effect sizes were quantified using the correlation coefficient *r*. To describe variability in these physical properties across speakers, we also computed speaker-by-speaker similarity matrices separately for each syllable and signal. For each pair of speakers, we calculated Pearson correlations between their time-aligned trajectories for the corresponding acoustic or facial-articulatory signal.

#### 2.4.2 Perceptual response on all stimuli across participants and speakers

To verify the perceptual properties of the dataset, we first calculated the proportions of each response choice (“ba,” “da,” “ga,” and “other”) to each stimulus. Given violations of normality, differences across syllables were tested using Friedman tests with Bonferroni-corrected Wilcoxon signed-rank tests for pairwise comparisons. Second, we ranked the speakers according to their McGurk illusion rate for each participant. Lastly, we assessed the consistency across participant in ranking the speakers according to their ability to elicit the illusion using Kendall’s coefficient of concordance (*W*).

#### 2.4.3 Predicting inter-participant variability of the McGurk illusion susceptibility

Following Dong et al. (2025), we first examined the associations between variability in susceptibility to the McGurk illusion and unisensory percepts using binomial generalized linear models (GLMs) with a logit link function (Equation 1). In the first model (Model 1), we predicted the probability (*pij*) of an illusory “da” response on McGurk trials from proportion of “ba” responses on A_ba_ trials (*X*_1*i*_, from now on auditory accuracy) and the proportion of “ga” responses on V_ga_ trials (*X*_2*i*_, from now on visual accuracy).

The significance of individual fixed-effect predictors was evaluated using Wald *z* tests, with two-way cluster-robust standard errors clustered by participant and speaker (Cameron & Miller, 2015). Multicollinearity was assessed using Variance Inflation Factors (VIF), with all predictors falling within acceptable limits (*VIF* < 5). We then extended the first model by incorporating predictors related to characteristic judgments of the McGurk stimuli. These predictors included the proportion of “congruent” responses on AV_m_ trials ( *X*_3*i*_,audiovisual congruency), the proportion of “synchronized” responses on AV_m_ trials (*X*_4*i*_, audiovisual synchrony), the mean rating of articulatory clarity on AV_m_ trials (*X*_5*i*_), the mean rating of perceived self-speaker similarity on AV_m_ trials (*X*_6*i*_,), and the mean rating of attractiveness on AV_m_ trials (*X*_7*i*_). Because audiovisual congruency and audiovisual synchrony exhibited substantial multicollinearity (*VIF* > 5), we fit two separate models (Model-2 and Model-3) with six predictors. Model-2 retained congruency and excluded synchrony, whereas Model-3 retained synchrony and excluded congruency. Predictors in both six-factor models were standardized to z scores prior to model fitting to facilitate comparison of regression coefficients.

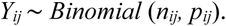

where

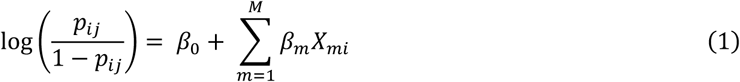

- *p*_*ij*_ : the probability of an illusory “da” response for participant *i* viewing speaker *j*.
- *β*_0_: the fixed intercept.
- *M*: the number of fixed-effect predictors; *M*=2 for Model-1 and *M*=6 for Models-2 and Models-3.
- *X*_*mi*_ : the m^th^ predictor for participant *i*. Across Models 1–3, predictors were coded as follows: auditory accuracy (*X*_1*i*_); visual accuracy (*X*_2*i*_); audiovisual congruency (*X*_3*i*_); audiovisual synchrony (*X*_4*i*_); articulatory clarity (*X*_5*i*_); self-speaker similarity (*X*_6*i*_); and attractiveness (*X*_7*i*_). Model-1 included *X*_1*i*_ and *X*_2*i*_ ; Model-2 included *X*_1*i*_, *X*_2*i*_, *X*_3*i*_, *X*_5*i*_, *X*_6*i*_, and *X*_7*i*_; and Model-3 replaced *X*_3*i*_ with *X*_4*i*_, while retaining the other predictors.
- *β*_m_: the fixed effects for predictor *X*_*mi*_.

Furthermore, we fitted Models 1-3 separately for each speaker to examine whether the associations between predictors (at participants-level) and the McGurk illusion rate were consistent across speakers. For each predictor, we calculated (i) the proportion of speakers for whom the predictor was significant after Benjamini-Hochberg false discovery rate (FDR) correction, with higher proportions indicating greater cross-speaker replicability; and (ii) the sign-congruency index (SCI), defined as (*N_pos_* - *N_neg_*) / *N_sig_*, where *N_pos_* and *N_neg_* denote the numbers of significant positive and negative coefficients, respectively, and *N_sig_* denotes the total number of significant coefficients. SCI values of 1 or -1 indicate that the direction of the fixed-effect coefficient (*β*) was consistent across all significant speaker-specific models.

#### 2.4.4 Predicting inter-speaker variability of the McGurk illusion susceptibility

To examine which variables predicted inter-speaker variability of the McGurk illusion susceptibility, we applied the same GLM analyses used in Section 2.4.3 to predict the probability (*p_ij_*) of the illusory “da” response on McGurk trials from speaker-level predictors (*X*_*j*_, averaged per speaker). The three models paralleled Models 1-3: the first included auditory accuracy (*X*_1*j*_) and visual accuracy (*X*_2*j*_) ; the second added audiovisual congruency (*X*_3*j*_), articulatory clarity (*X*_5*j*_), self-speaker similarity (*X*_6*j*_), and attractiveness (*X*_7*j*_). ; and the third replaced audiovisual congruency with audiovisual synchrony (*X*_4*j*_) while retaining the remaining predictors.

Similarly, we fitted each of the three models for every participant to examine whether the associations between predictors (at speaker-level) and the McGurk illusion rate were consistent across participants.

#### 2.4.5 Optimizing the McGurk illusion index

The large number of stimuli and behavioral responses enabled us to compare different variants of McGurk illusion indexes and to identify the optimal measure for future studies. Specifically, we assessed three indexes: (i) the sensory-weighted McGurk illusion index (SWMI), adapted from prior work (Stevenson et al., 2012); (ii) the normalized McGurk illusion index (NMI), adapted from prior studies (Gijbels et al., 2021; Grant et al., 1998); and (iii) the auditory-accuracy-scaled McGurk illusion index (AASMI), which quantifyied the proportional reduction of accuracy to the auditory /ba/ syllable after the incongruent visual /ga/ syllable was dubbed in McGurk stimuli.

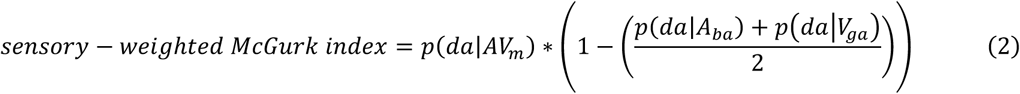

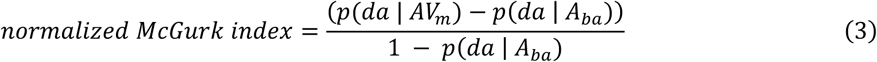

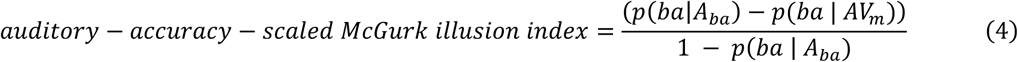

- *p* (da | AV_m_): the probability of a “da” response on McGurk trials.
- *p* (da | A_ba_): the probability of a “da” on auditory /ba/ trials.
- *p* (da | V_ga_): the probability of a “da” on visual /ga/ trials.
- *p* (ba | A_ba_): the probability of a “ba” on auditory /ba/ trials.
- *p* (ba | AV_m_): the probability of a “ba” response on McGurk trials.

A sensitive index reflecting the audiovisual integration should be separable from unisensory performance, particularly auditory confounds and be related to other audiovisual integration measures. We assessed these indices from three aspects. First, we calculated Pearson correlation coefficients among the indices. Second, we calculated Pearson correlation coefficients between each index and two benchmarks of audiovisual benefit: i. absolute gain, defined as the improvement from auditory to audiovisual performance; ii, and relative gain, defined as the proportion of the available improvement above auditory-only performance that was achieved when visual speech was added. For both sets of correlation analyses, *p* values were FDR-corrected. The computation of absolute and relative gain followed Magnotti et al. (2020) and Van Engen et al. (2017). Third, we tested the association between each index (dependent variable) and auditory accuracy (*X*_1*ij*_ ; independent variable), using the GLM at the participant level and speaker level separately.

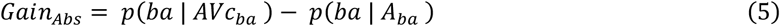

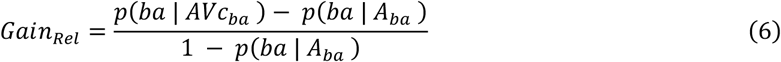

- *p* (ba | AVc_ba_): the probability of a “ba” response on congruent audiovisual /ba/ trials.
- *p* (ba | A_ba_): the probability of a “ba” response on auditory /ba/ trials; this index baseline auditory accuracy.
- *p* (ba | A_ba_): the remaining possible improvement over auditory performance.

## 3. Results

### 3.1 Differences in acoustic and visual articulatory signal across syllables and speakers

The acoustic spectra and facial articulations of the syllables /ba/, /da/, and /ga/ for a representative speaker (S048) are shown in Figure 2A.

**Fig. 2.**
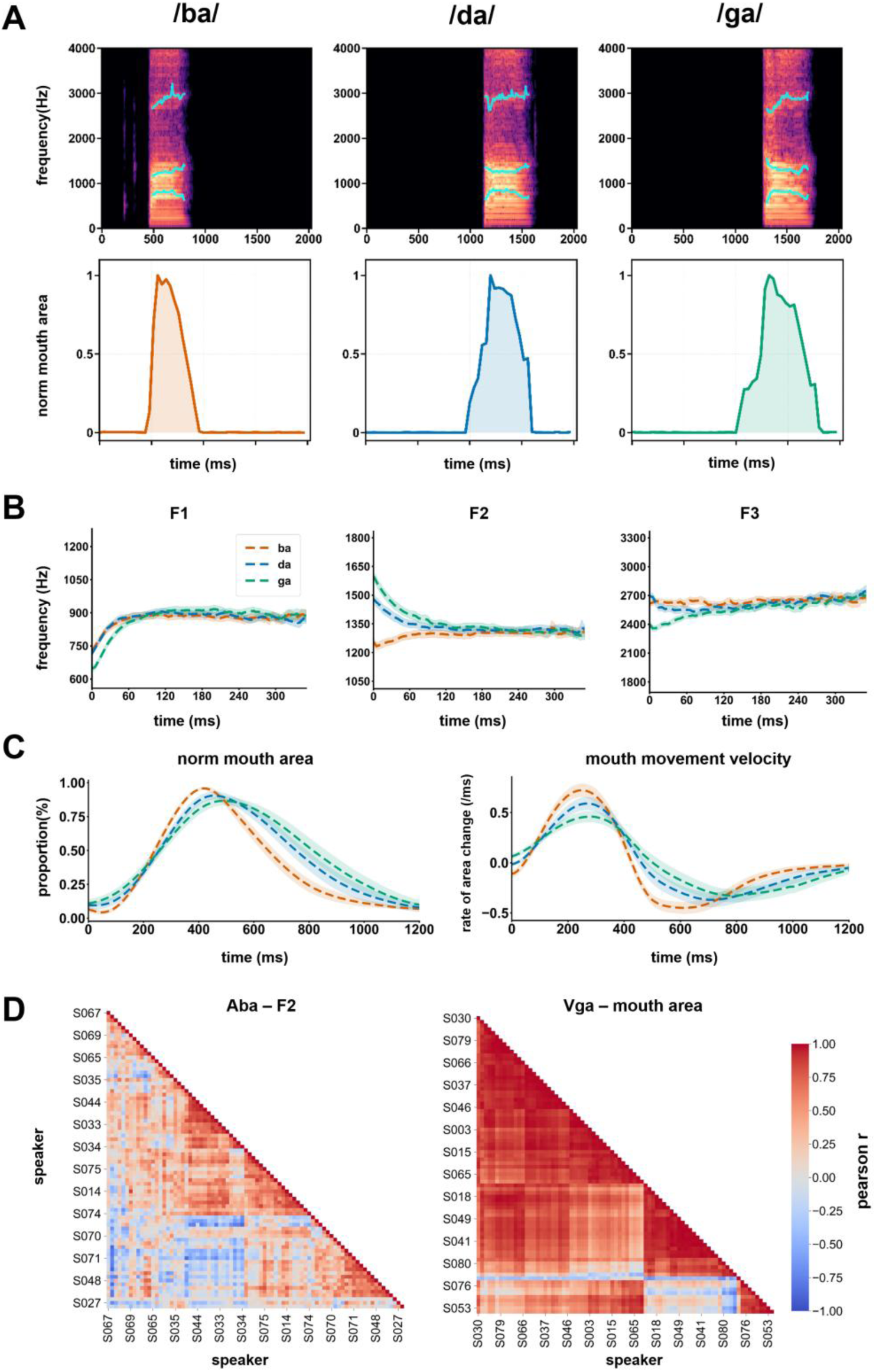
Physical properties of the McGurk stimuli. **(A)** Acoustic and visual articulatory signal of /ba/, /da/, and /ga/ syllables for a representative speaker (S048). The spectrograms with formant tracks (blue lines) illustrating the spectral evolution of F1, F2, and F3. **(B)** Mean formant trajectories of F1, F2, and F3, respectively, which aligned at voice onset (0 ms). Dashed lines represent the averaged frequency tracks for /ba/ (orange), /da/ (blue), and /ga/ (green), with shaded regions indicating 95% confidence intervals. **(C)** Mean visual articulatory kinematic trajectories aligned at the onset of mouth opening (0 ms), showing the temporal contours of mouth area and mouth movement velocity, respectively. **(D)** Pairwise speaker similarity matrices for the F2 trajectory of A_ba_ and the mouth-area trajectory of Vga. Rows and columns represent 80 speakers ordered by trajectory similarity.

For the onset formant frequencies (F1, F2 and F3), the Friedman test revealed a significant main effect of syllable type (A_ba_, A_da_, Aga) across speakers (Figure 2B; Table S1). Post-hoc comparisons indicated that the largest differences were observed between A_ba_ and A_ga_ (see Supplementary Results 1.1). Pearson correlations indicated that the similarity of acoustic trajectories varied across syllables and speakers (Figure S1), especially for the F2 trajectory of the A_ba_, the auditory counterpart of McGurk stimuli (Figure 2D).

For the articulation signal, a significant main effect of syllable type was observed across V_ba_, V_da_ and V_ga_ (Figure 2C; Table S1), with the largest differences again occurring between V_ba_ and V_ga_ (see Supplementary Results 1.1). The similarity of articulation trajectories varied across syllables and speakers (Figure S2), especially for the mouth-area trajectory of the V_ga_, the visual counterpart of McGurk stimuli (Figure 2D).

### 3.2 Response accuracy varied across syllables and modalities

For auditory trials, response accuracy differed significantly across target syllables (A_ba_, A_da_, A_ga_; χ²_(2)_ = 45.60, *p* < .001, Kendall’s *W* = .76; Figure 3A). Accuracy for A_ga_ was higher (*M* = 96%, *SD* = 6%) than for A_ba_ (*M* = 85%, *SD* = 9%*; p_bonf_ <* .001, *r =* .87) and A_da_ (*M* = 89%, *SD* = 8%; *p_bonf_ <* .001, *r =* .87), whereas accuracy for A_ba_ and A_da_ did not differ significantly.

**Fig. 3.**
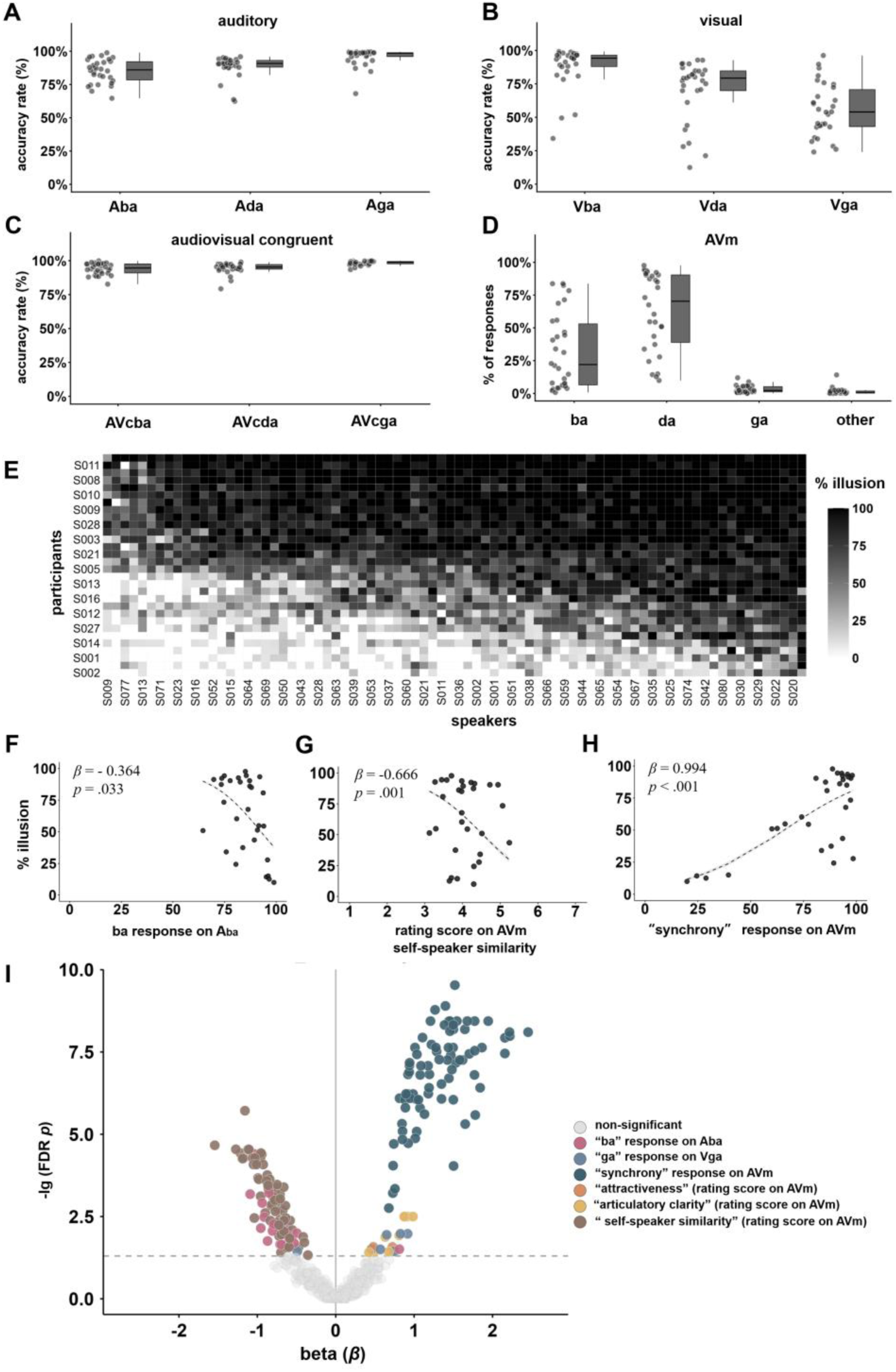
Perceptual responses to the McGurk stimulus and its counterparts. (A-C) Accuracy for auditory, visual, and audiovisual congruent syllables; each dot represents a participant; boxes show the interquartile range, center lines show medians, and whiskers extend to 1.5 times the interquartile range. (D) Response proportions to McGurk stimuli (AVm; auditory /ba/ paired with visual /ga/). (E) Heatmap of McGurk illusion rates for each participant-speaker pair, with rows and columns ordered by mean illusion rate. (F-H) Predicting participants’ McGurk illusion rates from audiovisual synchrony, auditory accuracy, and self-speaker similarity in Model 3, the model with the lowest AIC. Each dot represents a participant, and predictors were averaged across speakers. (I) Volcano plot of speaker-specific Model 3 estimates. Each dot is one fixed-effect estimate from a model fitted separately for one speaker; the x-axis shows the coefficient (*β*), and the y-axis shows −log10 (FDR-adjusted *p*), with higher values indicating stronger statistical significance. Colored dots indicate significant predictors, gray dots indicate non-significant predictors, vertical line marks *β* = 0, and the dashed horizontal line marks FDR-adjusted *p* = .05.

For visual trials, response accuracy differed significantly across target syllables (V_ba_, V_da_, V_ga_; χ²_(2)_ = 44.87, *p* < .001, Kendall’s *W* = .75; Figure 3B). Accuracy for V_ba_ (*M* = 88%, *SD* = 16%) was higher than for V_da_ (*M* = 71%, *SD* = 23%; *p_bonf_ <* .001, *r =* .84) and V_ga_ (*M* = 56%, *SD* = 20%; *p_bonf_ <* .001*, r =* .87), whereas accuracy for V_da_ and V_ga_ did not differ significantly.

For the audiovisual congruent trials, response accuracy differed significantly across the three congruent audiovisual syllables (AVc_ba_, AVc_da_, AVc_ga_; χ²_(2)_ = 36.60, *p* < .001, Kendall’s *W* = .61; Figure 3C), with higher accuracy for AVc_ga_ (*M* = 98%, *SD* = 2%) than for both AVc_ba_ (*M* = 94%, *SD* = 4%; *p_bonf_ <* .001*, r =* .85) and AVc_da_ (*M* = 94%, *SD* = 4%; *p_bonf_ <* .001*, r =* .86), whereas AVc_ba_ and AVc_da_ did not differ significantly.

These results parallel the patterns observed in the acoustic and facial articulatory signal. In the auditory modality, /da/ and /ba/ were more frequently misperceived than /ga/, whereas in the visual modality, /da/ and /ga/ were more frequently misperceived than /ba/. Full response distributions across the four response categories (“ba,” “da,” “ga,” and “other”) are provided in Supplementary Results and Figure S3.

### 3.3 McGurk illusion susceptibility varied across participants and speakers

For the McGurk trial, the proportions of the four responses options differed significantly (*χ²*_(3)_ = 74.78, Kendall’s *W* = 0.83, *p* < .001; Figure 3D). The proportions of illusory “da” (*M* = 63%, *SD* = 30%) was higher than that of “ba” response (auditory counterpart, *M* = 32%, *SD* = 29%; *p_bonf_* = .031*, r =* .51), which was higher than “ga” response (visual counterpart, *M* = 3%, *SD* = 3%; *p_bonf_* < .001*, r =* .83), which in turn exceeded “others” responses (*M* = 1%, SD = 3%; *p_bonf_* < .001, r = .71).

Susceptibility to the McGurk illusion varied substantially across participants (from 10% to 80%; Figure 3E) and speakers (from 33% to 90%). Participants who exhibited relatively low illusion rates for one speaker tended to do so for others, as indicated by consistent cross-speaker rankings (Kendall’s *W* = .728, χ²_(29)_ = 1690, *p* < .001).

### 3.4 Unisensory percepts and audiovisual characteristics predicted inter-participants variability in McGurk illusion susceptibility

In the first GLM model (Table S2), auditory accuracy negatively predicted illusion rates (*β* = −0.702, *SE* = 0.334, *z* = -2.10, *p* = .036), replicating the finding that higher auditory precision is associated with reduced McGurk susceptibility (Dong et al., 2025).

In the second GLM model (Table S3), audiovisual congruency positively predicted illusion rates (*β* = 0.577, *SE* = 0.188, *z* = 3.07, *p* = .002), indicating that an increase in perceived congruency was associated with higher illusion rates. Self-speaker similarity negatively predicted illusion rates (*β* = -0.535, *SE* = 0.248, *z* = -2.16, *p* = .031), indicating that an increase in perceived self-speaker similarity was associated with lower illusion rates.

In the third GLM model (Table S4), auditory accuracy (*β* = -0.364, *SE* = 0.171, *z* = -2.13, *p* = .033; Figure 3F) and self-speaker similarity (*β* = -0.666, *SE* = 0.204, *z* = -3.26, *p* = .001; Figure 3G) negatively predicted illusion rates, and audiovisual synchrony positively predicted illusion rates (*β* = 0.994, *SE* = 0.150, *z* = 6.62, *p* < .001; Figure 3H). Visual accuracy, attractiveness, and articulatory clarity did not reach significance in either extended model (all *ps* > .130). These results suggest that the variabilities of McGurk illusion rate across participants were associated not only with differences in unisensory perception but also with variations in audiovisual correspondence (i.e., congruency and synchrony) and higher-level characteristics (self-speaker similarity).

To assess whether these prediction effects from participants’ perspective were consistent across speakers, we fit the model with lowest AIC (i.e., the third GLM model: 14,281.44) separately for each of the 80 speakers (Figure S4). In this model (Figure 3I), audiovisual synchrony (100%; SCI = 1.0) and self-speaker similarity (81.3%; SCI = −1.0) were significant for a majority of speakers, with consistent coefficient directions. Auditory accuracy (26.3%; SCI = −0.9) and articulatory clarity (12.5%; SCI = 1.0) were significant for fewer speakers, also with consistent coefficient directions. Attractiveness (2.5%) and visual accuracy (7.5%) were significant for few speakers.

### 3.5 Unisensory percepts and audiovisual characteristics predicted inter-speakers variability in McGurk illusion susceptibility

In the first model (Table S5), both auditory accuracy (*β* = -0.33, *SE* = 0.07, *z* = -4.83, *p* < .001) and visual accuracy (*β* = -0.14, *SE* = 0.05, *z* = -2.91, *p* = .004) negatively predicted illusion rates, indicating that higher unisensory precision is associated with reduced McGurk susceptibility (Dong et al., 2025).

In the second model (Table S6), in addition to the negative effect of auditory accuracy (*β* = -0.244, *SE* = 0.057, *z* = -4.28, *p* < .001; Figure 4A), audiovisual congruency positively predicted illusion rates (*β* = 0.218, *SE* = 0.061, *z* = 3.56, *p* < .001; Figure 4B), indicating that McGurk stimuli perceived as more congruent elicited higher illusion rates. Articulatory clarity also positively predicted illusion rates (*β* = 0.17, *SE* = 0.054, *z* = 3.17, *p* = .002; Figure 4C), suggesting that if the perceivers thought that the articulation in McGurk stimulus was clearer, the illusion rate on that stimulus was higher. Visual accuracy, attractiveness, and self-speaker similarity were not significant (all *ps* > .121).

**Fig. 4.**
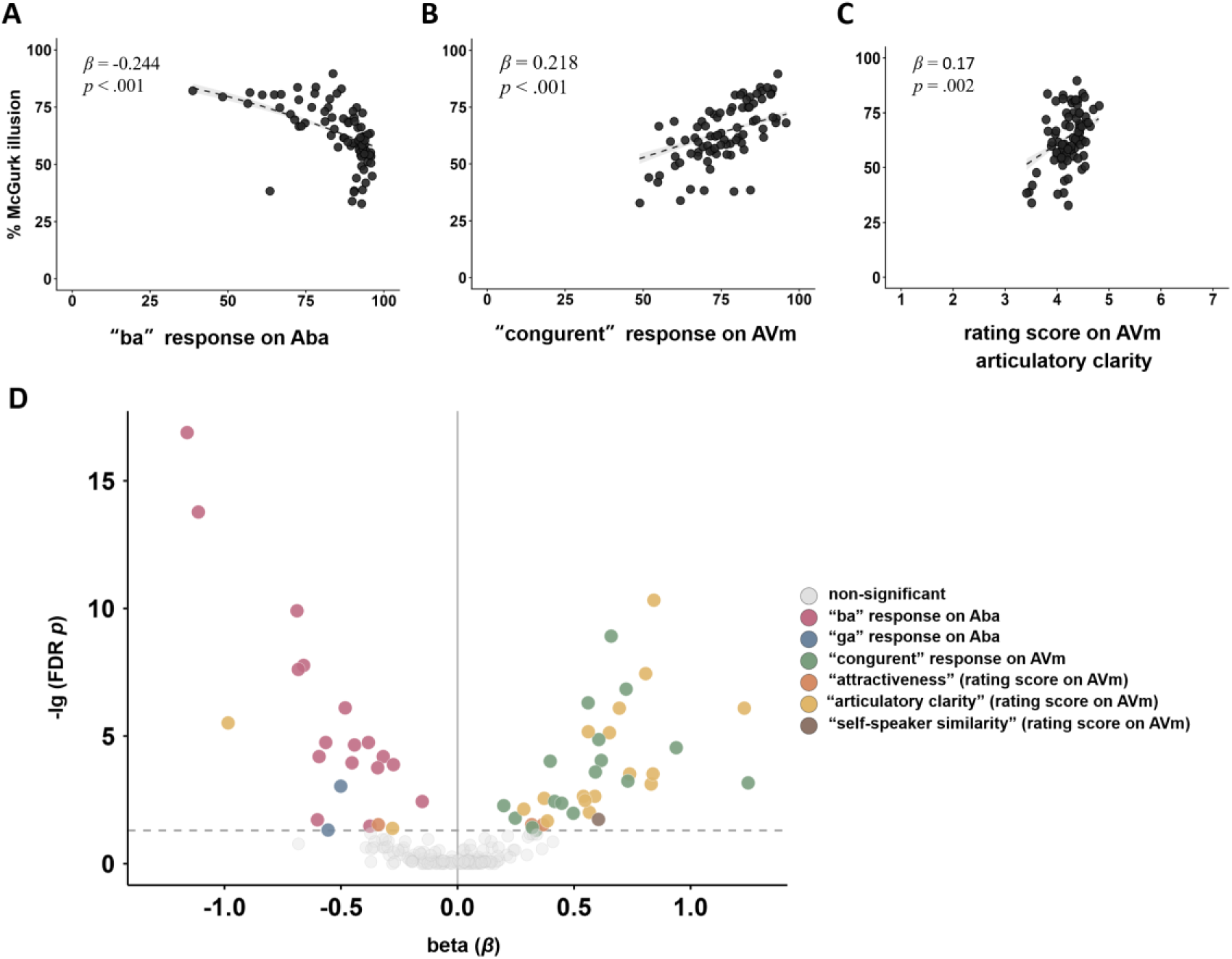
Predicting of McGurk illusion susceptibility across speakers. (A-C) Predicting McGurk illusion rates on speakers from audiovisual congruency, auditory accuracy, and articulatory clarity in Model 2, the model with the lowest AIC. Each dot represents one speaker, and predictors were averaged across participants; dashed lines and shaded bands show fitted logistic curves and 95% confidence intervals. (D) Volcano plot of participant-specific Model 2 estimates. Each dot is one fixed-effect estimate from a model fitted separately for one participant; the x-axis shows the coefficient (*β*), and the y-axis shows −log10 (FDR-adjusted *p*), with higher values indicating stronger statistical significance. Colored dots indicate FDR-significant predictors, gray dots indicate non-significant predictors, the vertical line marks *β* = 0, and the dashed horizontal line marks FDR-adjusted *p* = .05.

In the third model (Table S7), the effects of auditory accuracy (*β* = -0.28, *SE* = 0.061, *z* = -4.61, *p* < .001) and articulatory clarity (*β* = 0.227, *SE* = 0.056, *z* = 4.06, *p* < .001) remained significant. In parallel to the audiovisual congruency, audiovisual synchrony positively predicted illusion rates (*β* = 0.186, *SE* = 0.052, *z* = 3.59, *p* < .001). These results indicate that the variabilities in McGurk illusion rate across speakers were influenced by variations in unisensory perception of the perceivers, differences in articulation features of the speakers, and variations in audiovisual correspondence of the audiovisual stimuli.

Furthermore, we fitted the models with the lowest AIC (i.e., the second model: 18,906.86) separately fitted for each participant (Figure S5). In this model (Figure 4D), articulatory clarity (60.0%; SCI = 0.78), audiovisual congruency (56.7%; SCI = 1.0), and auditory accuracy (56.7%; SCI = −1.0) were significant for a majority of participants. Attractiveness (10.0%), visual accuracy (6.7%), and self-speaker similarity (3.3%) were significant for only a small proportion of participants.

### 3.6 Candidate McGurk illusion indices showed complementary trade-offs

All three candidate indices exhibited substantial variation (Figure 5A-B, Table S8) and were significantly correlated across participants (all *rs* ≥ .93; Figure 5C) and speakers (all *rs* ≥ .63; Figure 5D).

**Fig. 5.**
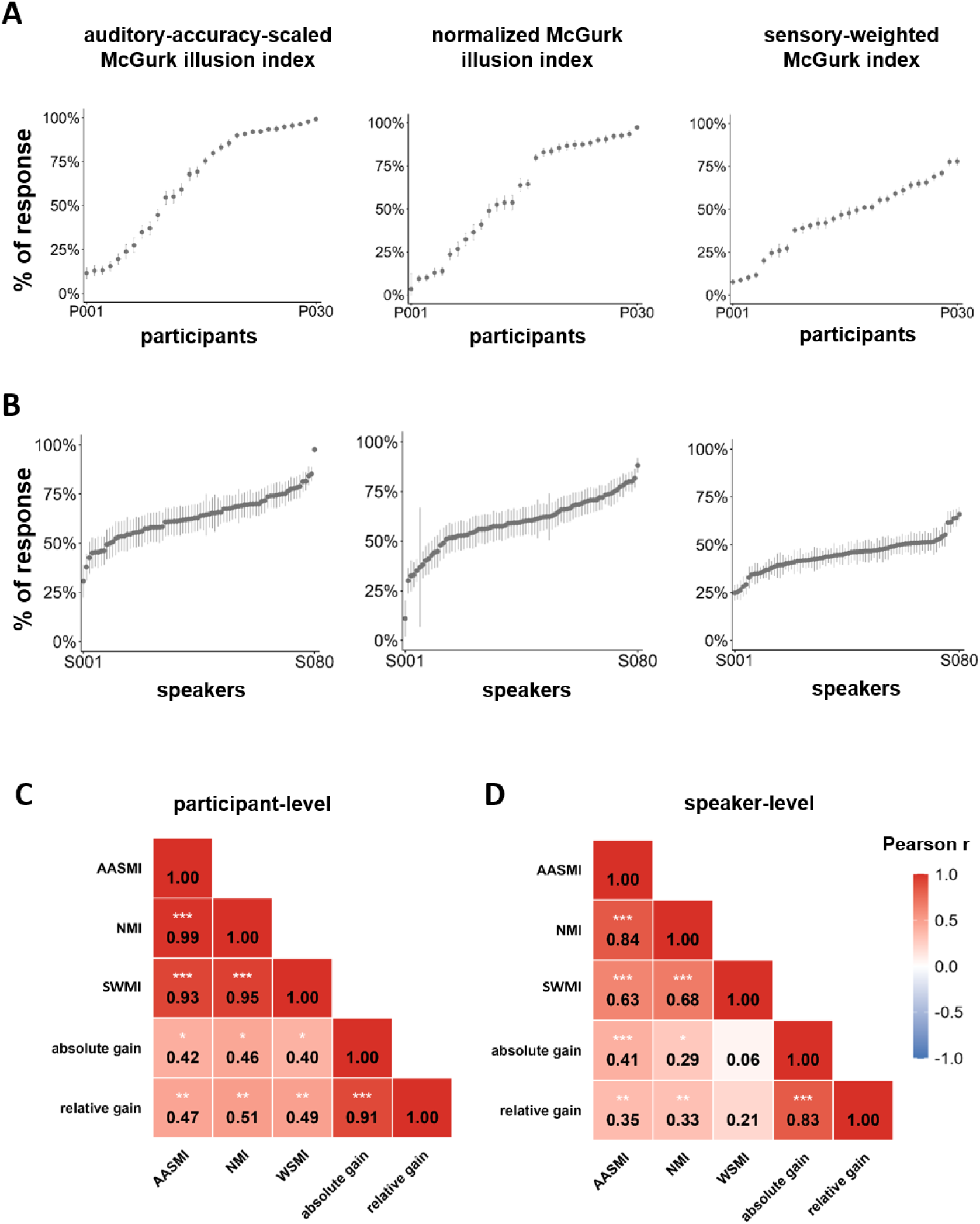
Variations and associations of candidate McGurk illusion indices. (A) Participant-level ranked distributions of the auditory-accuracy-scaled McGurk illusion index, the normalized McGurk illusion index, and the sensory-weighted McGurk index. Each point represents one participant. Rank labels P001 and P030 denote the lowest and highest ranked positions, respectively. (B) Speaker-level ranked distributions of the same three indices. Each point represents one speaker. (C) Pearson correlations among the three candidate indices, absolute gain and relative gain across participants. (D) Pearson correlations among the same variables across speakers. AASMI = auditory-accuracy-scaled McGurk illusion index; NMI = normalized McGurk index; SWMI = sensory-weighted McGurk index. \**p* < .05; \*\**p* < .01; \*\*\**p* < .001.

Across participants, the auditory-accuracy-scaled McGurk illusion index ranged from 12% to 99%, the normalized McGurk illusion index ranged from 3% to 97%, and the sensory-weighted McGurk illusion index ranged from 8% to 78% (Figure 5A). Auditory accuracy did not significantly predict the auditory-accuracy-scaled McGurk illusion index (*b* = -0.11, *p* = .057) or the sensory-weighted McGurk illusion index (*b* = -0.06, *p* = .093), but significantly predicted the normalized McGurk illusion index (*b* = -0.12, *p* = .033; Table S9). All three candidate indices were positively correlated with both relative gain (*r* = .47–.51, *ps* < .01) and absolute gain (*r* = .40–.46, *ps* < .05; Figure 5C).

Across speakers, the auditory-accuracy-scaled McGurk illusion index ranged from 31% to 98%, the normalized McGurk illusion index ranged from 11% to 88%, and the sensory-weighted McGurk illusion index ranged from 25% to 66% (Figure 5B). Auditory accuracy predicted the auditory-accuracy-scaled McGurk illusion index (*b* = - 0.05, *p* < .001), but not the normalized McGurk illusion index (*b* = -0.03, *p* = .077) or the sensory-weighted McGurk illusion index (*b* = 0.003, *p* = .739; Table S9). The auditory-accuracy-scaled McGurk illusion index (absolute gain: *r* = .41, *p* < .001; relative gain: *r* = .35, *p* < .01) and the normalized McGurk illusion index (absolute gain: *r* = .29, *p* < .05; relative gain: *r* = .33, *p* < .01) were significantly correlated with both absolute and relative gain. The sensory-weighted McGurk illusion index was not significantly correlated with either absolute gain (*r* = .06, *p* > .05) or relative gain (*r* = .21, *p* > .05; Figure 5D).

## Discussion

We introduced the largest McGurk illusion dataset (MID) to date, with one thousand four hundred and forty high quality stimuli, for audiovisual integration and speech perception research. We conducted rigorous unisensory and audiovisual laboratory tasks to examine the physical and perceptual properties of the McGurk stimuli. The results indicated that, at physical level, McGurk illusion syllables (i.e., /ba/, /da/, and /ga/) differed in acoustic and facial articulatory features across syllables and speakers. The syllable /da/ was closer to /ba/ in acoustic dimension, closer to /ga/ in articulation dimension, and these features exhibited substantial variation across speakers. At the perceptual level, the McGurk susceptibility varied across participants and speakers. These variabilities were associated with not only variations in participants’ unisensory precision but also differences in audiovisual correspondence (e.g., audiovisual congruency and synchrony) and speakers characteristics (e.g., self-speaker similarity). In the comparison of McGurk illusion-based indices, the sensory-weighted and auditory-accuracy-scaled McGurk illusion indices performed better at the participant level, whereas the normalized McGurk illusion index performed better at the speaker level.

Using this stimulus set, we observed substantial differences in acoustic and facial articulatory features across McGurk illusion syllables and speakers. The illusory syllable /da/ was closer to syllable /ba/ in acoustic features but closer to syllable /ga/ in facial articulation features. Both acoustic signal and facial articulatory measures varied across speakers. These physical differences reflect well-established cues to speech categorization (Bent & Holt, 2017; Jiang & Bernstein, 2011; Johnson & Sjerps, 2021; Olmstead et al., 2020). For the acoustic feature, formant transitions, particularly F2 and F3, provide important cues to place-of-articulation contrasts among stop consonants such as /ba/, /da/, and /ga/ (Blumstein & Stevens, 1979; Kewley-Port, 1983). For the articulation feature, McGurk illusion syllables differ in the facial visibility of articulatory gestures, with bilabial /ba/ providing more salient lip-based information than the more posterior articulations involved in /da/ and /ga/ (Chartier et al., 2018). From a Bayesian cue-integration perspective, differences in these physical features across speakers may influence McGurk susceptibility by changing the reliability and informativeness of the auditory and visual signals (Magnotti & Beauchamp, 2017; Meijer & Noppeney, 2023).

We examined the validity of the stimulus set by replicating key findings from previous McGurk illusion studies. Our stimuli elicited a robust McGurk illusion, with a mean fusion rate of 63%, comparable to previous study from Chinese speakers (Dong et al., 2025) and to non-Chinese speakers (Basu Mallick et al., 2015; Magnotti et al., 2025; Stropahl et al., 2017). The illusion rates ranged from 10% to 80% across participants, and from 33% to 90% across speakers, which were consistent with previous studies (Basu Mallick et al., 2015; Brown et al., 2018; Strand et al., 2014). The MID also showed that the tendency of particular speakers to elicit higher illusion rates was relatively consistent across participants, also replicating our previous findings (Dong et al., 2025). Moreover, our results characterized multiple key sources of McGurk illusion variability across participants and speakers. At both the participant and speaker levels (Dong et al., 2025), higher unisensory perceptual precision was associated with lower illusion susceptibility, whereas higher perceived audiovisual congruency and synchrony were associated with higher illusion susceptibility. These findings are consistent with prior evidence that McGurk illusion depends on temporal alignment between auditory and visual speech (Van Wassenhove et al., 2007), and that perceived audiovisual correspondence or disparity can influence whether conflicting speech cues are fused or segregated (Magnotti & Beauchamp, 2017). Furthermore, our results identified the effects of speaker-related characteristics. Higher self-speaker similarity was associated with lower illusion rates in participants, and higher articulatory clarity was associated with higher illusion susceptibility across speakers. These findings suggest that variation in speaker familiarity and the intelligibility of a speaker’s articulation influence the audiovisual speech processing (Alsius et al., 2018; Aruffo & Shore, 2012; Tye-Murray et al., 2013).

Further analyses with models fitted separately for each participant and each speaker showed that each factor had relatively consistent influences on McGurk illusion susceptibility, while the statistical detectability of these influences depended on the particular participants and speakers included. These results may explain why studies using smaller or different stimulus sets have reported inconsistent associations between McGurk susceptibility, unisensory speech perception, and other measures of audiovisual speech perception (Brown et al., 2018; Strand et al., 2014; Van Engen et al., 2017). Together, these findings underscore the need to characterize participant-specific and speaker-specific contributions within a common stimulus framework (Magnotti & Beauchamp, 2015).

The candidate McGurk illusion indices showed different trade-offs at the participant and speaker levels. The sensory-weighted and auditory-accuracy-scaled McGurk illusion indices reduced dependence on auditory accuracy while retaining associations with audiovisual gain at the participant level; at the speaker level, the normalized McGurk illusion index achieved the clearest balance between these two criteria. The sensory-weighted index downweights /da/ responses that also occur in the auditory-only /ba/ and visual-only /ga/ conditions, reducing responses attributable to unisensory misidentification rather than the influence of visual /ga/ (Stevenson et al., 2012); this correction has been shown to improve sensitivity in clinical group comparisons (Butera et al., 2023). The auditory-accuracy-scaled index captures the proportional reduction in correct /ba/ identification when incongruent visual /ga/ is added. The normalized McGurk illusion index subtracts the auditory-only /ba/-to-/da/ baseline and scales the residual—a baseline-corrected normalization analogous to audiovisual gain computation (Gijbels et al., 2021; Grant et al., 1998). These trade-offs suggest that these indices reduce specific sources of confounding, but may not isolate audiovisual integration per se. Future work may benefit from model-based approaches, such as Bayesian causal-inference models, that estimate sensory precision, causal priors, and integration-related parameters separately (Zhu et al., 2026).

Beyond its use as a standardized resource for McGurk illusion research, our data set may support several lines of research. First, in the speaker normalization research, it provides a broad and controlled acoustic space, with standardized stimuli from a large set of speakers, for examining how listeners map talker-specific acoustic variation onto stable phonetic categories (Bhaya-Grossman & Chang, 2022; Johnson, 2019; Oganian et al., 2023). Second, the MID can support research on speaker identity and speaker variability, including how listeners represent facial and vocal identity cues across speakers and utterances (Lally et al., 2023; Lavan & Sutherland, 2024). Third, the MID provides a controlled benchmark for testing how artificial neural networks use visible articulatory cues, resolve audiovisual conflict, and generalize across speakers. It also supports comparisons of audiovisual integration mechanisms between human participants and multimodal models (Grasse & Tata, 2025; Ma et al., 2026).

While our dataset exhibited substantial validity, several limitations should be acknowledged. First, the MID focused on the canonical McGurk illusion syllables (i.e., auditory /ba/ with visual /ga/), which improved comparability with prior work but limits phonetic coverage. Future research could include additional consonants, vowels, and audiovisual pairings. Second, both speakers and participants were Chinese speaking adults. Given the influences of developmental, aging, and cultural on speech perception, broader sampling across age, ethnic, and language backgrounds will be needed to improve generalizability of the dataset (Alsius et al., 2018; Gijbels et al., 2021; Magnotti et al., 2025; Sekiyama et al., 2014). Third, the validation focused mainly on McGurk illusion measures, leaving broader audiovisual speech integration effects, such as sentence recognition in noise (Magnotti et al., 2020; Van Engen et al., 2017). Future work could combine the MID with broader audiovisual speech measures to clarify how McGurk susceptibility relates to audiovisual speech perception in naturalistic communication contexts.

In conclusion, we introduce the McGurk illusion dataset (MID), which is the largest publicly available McGurk stimulus dataset to date. This dataset comprises one thousand four hundred and forty high quality stimuli from eighty speakers and was validated through rigorous behavioral judgments. Using MID, we revealed the large variations of acoustic and articulatory features of McGurk illusion syllables, replicated substantial inter-participant and inter-speaker variabilities in illusion susceptibility, and demonstrated associations between illusion susceptibility and other audiovisual processes. Furthermore, the stimulus set enabled systematic comparisons of the reliability of different McGurk illusion-based indices of audiovisual speech integration. Overall, the MID not only provides a standardized resource for investigating audiovisual speech integration and its alterations across populations, but also supports research on speaker normalization, lip-reading, and speech perception.

## Declarations

### Ethics approval

This study was approved by the Ethics Committee of the School of Psychology at South China Normal University (SCNU-PSY-2024-435).

### Consent to participate

All participants provided written informed consent and received monetary compensation.

### Consent for publication

All participants were informed that their anonymized video, voice, and behavior data would be stored in a publicly accessible archive provided by South China Normal University and used in future research projects.

### Open Practices Statement

All stimuli and scripts will be made publicly available on an official website for non-commercial use upon publication of the manuscript, subject to the consent and data-usage conditions approved by the Ethics Committee of the School of Psychology at South China Normal University.

### Authors’ contributions

Zhengye Wang, conceptualization, stimulus collection, data collection, data analysis, writing manuscript, revising manuscript. Gantang Li, writing manuscript, revising manuscript. Yating Yu, Junyi Wu, and Zenghui Yu, stimulus collection, generating stimuli, data collection. Yang Meng, resources. Suiping Wang, revising manuscript, supervision, resources and funding acquisition. Chenjie Dong, conceptualization, writing manuscript, revising manuscript, supervision, and funding acquisition.

## Acknowledgments

The authors would like to thank all the speakers and perceivers participating in this study.

